# Increased Risk for Cerebral Small Vessel Disease is Associated with Quantitative Susceptibility Mapping in HIV Infected and Uninfected Individuals

**DOI:** 10.1101/2021.03.13.435243

**Authors:** Kyle D. Murray, Md Nasir Uddin, Madalina E. Tivarus, Bogachan Sahin, Henry Z. Wang, Meera V. Singh, Xing Qiu, Lu Wang, Pascal Spincemaille, Yi Wang, Sanjay B. Maggirwar, Jianhui Zhong, Giovanni Schifitto

## Abstract

The aim of this study was to assess in the context of cerebral small vessel disease (CSVD), cardiovascular risk factors and white matter hyperintensities (WMHs) were associated with brain tissue susceptibility as measured by quantitative susceptibility mapping (QSM). Given that CSVD is diagnosed by the presence of lacunar strokes, periventricular and deep WMHs, increased perivascular spaces, and microbleeds, we expected that QSM could capture changes in brain tissue due to underlying CSVD pathology. We compared a cohort of 101 HIV-infected individuals (mean age (SD) = 53.2 (10.9) years) with mild to moderate cardiovascular risk scores, as measured by the Reynold’s risk score, to 102 age-matched controls (mean age (SD) = 50.3 (15.7) years) with similar Reynold scores. We performed brain MRI to assess CSVD burden by acquiring 3D T1-MPRAGE, 3D FLAIR, 2D T2-TSE, and mGRE for QSM. We found that signs of CSVD are significantly higher in individuals with HIV-infection compared to controls and that WMH volumes are significantly correlated with age and cardiovascular risk scores. Regional QSM was associated with cardiovascular risk factors, age, sex, and WMH volumes but not HIV status. These results suggest that QSM may be an early imaging marker reflective of alterations in brain microcirculation.

## 1 INTRODUCTION

Cerebrovascular disease (CBVD), affecting both large and small vessels, is among the highest leading causes of death in the United States, resulting in 45.2 deaths per 100,000 people in 2018 (Center for Health Statistics, 2018). In this study, we focused on cerebral small vessel disease (CSVD), which is common in older adults, causes ~25% of all ischemic strokes worldwide (Debette & Markus, 2010; ter Telgte et al., 2018; Wardlaw et al., 2013), and is a leading cause of cognitive impairment and dementia (Horsburgh et al., 2018). CSVD is a disease of small blood vessels and can be diagnosed by magnetic resonance imaging (MRI) by detecting lacunae, white matter hyperintensities (WMHs), hemorrhagic strokes, brain atrophy, and enlarged perivascular spaces (Wardlaw et al., 2013). CSVD pathogenesis is not fully understood and is likely multifactorial. Several factors are thought to contribute, including neuroinflammation, vessel stiffness, and altered blood brain barrier (BBB) (Aribisala et al., 2014; Wardlaw et al., 2013, 2017).

HIV-infection tends to display accelerated signs of CSVD, possibly due to persistent chronic immune activation, which may affect intrinsic brain properties such as tissue microstructure (Hearps et al., 2012; Martin et al., 2013). HIV impacts the cardiovascular system and can lead to increased risk of cardiovascular disease (CVD) and CBVD, despite effective combination antiretroviral therapies (cART) that controls viremia. Furthermore, there is evidence that cART itself can contribute to CBVD (Edwards et al., 2016; Singh et al., 2019). As a result of cART treatment, HIV-infected (HIV+) individuals are able to live much longer lives, making them more susceptible to developing common neurodegenerative diseases that affect the general population (Vinikoor et al., 2013). While many studies have focused on HIV-associated CBVD, much less attention has been devoted to HIV-associated CSVD, in spite of studies showing that HIV and CBVD share certain cardiovascular risk factors, such as diabetes and hypertension (HTN) (Vinikoor et al., 2013). Additionally, these cardiovascular risk factors are known to be associated with chronic vascular inflammation (Lopez-Candales et al., 2017).

CSVD pathology is expected to alter brain iron distribution and myelin integrity. Excessive iron accumulation in the subcortical deep gray matter (DGM) is implicated in several neurodegenerative diseases predominantly impacting the white matter (Thomas & Jankovic, 2004). It has been reported that iron accumulation in the DGM may contribute to neurodegeneration through the generation of reactive oxygen species (Farina et al., 2013; Yan et al., 2013). To date, several MRI techniques have been developed to estimate iron content in the brain, including transverse relaxation mappings (R2, R2*, and R2’), susceptibility weighted imaging (SWI), field dependent rate increase (FDRI), and quantitative susceptibility mapping (QSM) (Deistung et al., 2013; Stankiewicz et al., 2007; Uddin et al., 2016). Among these techniques, QSM is considered to be the most sensitive and reliable method to study iron in the brain and can be used to assess brain tissue susceptibility in pathologies (de Rochefort et al., 2010; Du et al., 2016; Haacke et al., 2015; Langkammer et al., 2016; Wang & Liu, 2015). QSM also appears to have lower variability and higher repeatability to quantify iron compared to R2* mapping (Feng et al., 2018). Further, the magnetic susceptibility provided by QSM reflects the paramagnetic (iron) and diamagnetic (myelin and brain parenchyma) contributions of susceptibility sources in brain tissue, while R2* mapping shows similar effects for both types of susceptibility. That is, QSM is sensitive to non-heme iron deposition in the iron rich DGM and demyelination in the white matter which may be linked to WMHs and other indicators of ischemia and cerebral microbleeds, all of which are clinical markers for CSVD (Yan et al., 2013). Limited information in the literature is available on changes in brain tissue susceptibility due to CVD.

In this study, we hypothesized that increased cardiovascular risk was associated with altered microcirculation and damage to the BBB, leading to changes in brain iron deposition, thus altering tissue susceptibility properties as measured by QSM. Therefore, the aims of our study were: to investigate the non-heme iron related changes in the presence of HIV-infection and CSVD using QSM and R2*mapping and to investigate the association between iron deposition in the DGM and cardiovascular risk factors.

## 2 MATERIALS AND METHODS

### 2.1 Research Subjects

Study procedures are fully described in Murray et al. (2020). As part of an ongoing study approved by the Research Subjects Review Board (RSRB) at the University of Rochester, one hundred one HIV+ subjects (mean ± SD age = 53.2 ± 10.9 years) were evaluated to study the effects of CSVD in HIV patients. One hundred two HIV– uninfected (HIV−) controls (mean ± SD age = 50.3 ± 16.7 years) were recruited and matched for age and sex, as possible. Neurocognitive and functional performance assessments were performed on all subjects as part of the study’s screening procedures. Written informed consent was obtained prior to any evaluation according to RSRB approval.

In order to evaluate the effects of changes in tissue susceptibility in HIV-infection and CSVD, we split our total dataset into four cohorts based on HIV- and CSVD-status (e.g., Healthy Controls (HIV−CSVD−); Controls with CSVD (HIV−CSVD+); HIV-infected without CSVD (HIV+CSVD−); and HIV-infected with CSVD (HIV+CSVD+)). CSVD+ indicates a Fazekas score reading of greater than 0 for deep or periventricular readings. WMH burden was assessed by Fazekas scores for deep and periventricular WMHs (Fazekas et al., 1987). Due to the lack of high CSVD burden in our dataset, we do not stratify CSVD status by each Fazekas rating (1 – 3).

Variables of interest included in these analyses include HIV markers, demographic information, clinical covariates, and MRI. HIV markers include CD4+ count, viral load (VL), medication regimen, and cerebrovascular risk factors that contribute to the Reynold’s risk score (RRS) (i.e., sex, age, smoking status, systolic blood pressure, total cholesterol, HDL cholesterol, high sensitivity C-reactive protein (hsCRP), and immediate family stroke history) (Ridker et al., 2007). Demographic variables considered as covariates were age, sex, and relevant prior cerebrovascular medical history, including diagnoses of hypertension (HTN) and diabetes type II (DIA). MRI was used to derive QSM and R2* maps for whole brain and region-of-interest (ROI) calculations and qualitative diagnoses of CSVD via deep and periventricular Fazekas scores.

### 2.2 MRI Acquisition and Processing

All imaging was conducted on a research dedicated 3T whole-body Siemens MAGNETOM Prisma scanner (Erlengen, Germany) equipped with a 64-channel head coil transmission. High-resolution T1-weighted (T1w) anatomical images were acquired using the magnetization prepared 3D rapid gradient echo (MPRAGE) sequence (inversion time (TI)= 926 ms; repetition time (TR) = 1840.0 ms; echo time (TE) = 2.45 ms; flip angle = 8°; echo spacing (ESP) = 7.5 ms; receiver bandwidth (RBW) = 190 Hz/pixel; resolution = 1 mm isotropic). QSM were acquired with a 3D multi-gradient echo (mGRE) pulse sequence with bi-polar readout (TR = 84 ms; TE (1^st^ echo) = 5.43 ms; number of echoes =8; ESP = 1.5 ms; RBW = 930 Hz/pixel; GRAPPA = 2; resolution = 0.94 × 0.94 × 2.0 mm ^3^; scan time = 7:7 min). In addition to the T1w and QSM sequences that were used quantitatively, a 2D T2-weighted (T2w) turbo spin echo sequence (TR = 6000 ms; TE = 100 ms; ESP = 11.1 ms; flip angle = 150°; rBW = 222 Hz/pixel; resolution = 0.5 × 0.5 × 5 mm^3^; scan time = 2:54 min) and 3D fluid attenuated inversion recovery (FLAIR) sequence (TI=1800 ms; TR = 5000 ms; TE = 100 ms; ESP = 3.42 ms; GRAPPA = 2; rBW = 751 Hz/pixel; resolution = 1 mm isotropic; scan time = 5:40 min) were acquired for diagnostic readings. The team’s radiologist reviewed the T1w, T2w, FLAIR, and T2* (i.e., 1/R2*) images to qualitatively assess CSVD burden for each subject. WMH burden was assessed via the FLAIR images. Additionally, VolBrain was used to quantitatively measure the total WMH burden using the T1w and FLAIR images (Manjón & Coupé, 2016).

Structural segmentation was performed on the T1w images using FSL (Jenkinson et al., 2012; Woolrich et al., 2009). The processing pipeline includes image reorientation and cropping, radio-frequency bias-field correction, linear and nonlinear registration to Montreal Neurological Institute (MNI)-152 2mm standard space, brain extraction, tissue segmentation, and subcortical structure segmentation. Regions of interest (ROIs) considered in this study include regions previously linked to susceptibility changes reflecting underlying pathology. These ROIs include the Amygdala (Amyg), Caudate (Cau), Globus Pallidus (GPal), Hippocampus (Hip), Nucleus Accumbens (Nac), Putamen (Put), Thalamus (Tha), global gray (GM) and white matter (WM), and global whole brain matter (WHB). Voxel-based whole-brain analyses (VBA) were used to determine any regions of interest in addition to the regions described above.

QSM reconstruction was performed utilizing the MEDI+0 toolbox (de Rochefort et al., 2010; Yao et al., 2017), on the MATLAB R2018a environment (The Mathworks, Inc., Natick, MA). Phase unwrapping was performed and background field removal was completed using the projection onto dipole field (PDF) method. R2* maps were produced via mono-exponential fitting of the magnitude images and used to create cerebrospinal fluid (CSF) masks as anatomical prior information before performing MEDI. Finally, the dipole inversion was performed using L1 regularization with Lagrange multipliers λ set to 1000 and λ_CSF_ set to 100, spherical mean value (SMV) radius set to 5, and using the model error reduction through iterative tuning (MERIT) option (J. Liu et al., 2012; T. Liu et al., 2011, 2013).

### 2.3 Statistical Analysis

For the region-based analysis, native space imaging maps were warped into MNI152-2mm space using FLIRT and FNIRT (J L R Andersson et al., 2010; Jesper L R Andersson et al., 2007). The ROIs were then applied as masks to the images from the Harvard-Oxford cortical and subcortical atlases. The mean metric value and standard deviation were determined using FSLSTATS for each ROI. Two-way analysis of variance (ANOVA) analyses were performed for each ROI to assess QSM and R2* differences between the four different cohorts, followed by post-hoc t-tests between each group pair. For the whole-brain analysis, images were warped into standard space using FNIRT. Gaussian smoothing of 5mm was performed to correct for coregistration errors and other susceptibility induced imperfections. An average template was calculated from all smoothed images. Randomise was used with 5000 permutations to perform the whole-brain QSM statistical analysis comparing HIV+ subjects to HIV- controls (Winkler et al., 2014).

We performed Pearson correlation analyses to study the marginal associations between imaging metrics and age and RRS, and between deep white matter hyperintensities (DWMHs) and periventricular white matter hyperintensities (PVWMHs). Multivariate linear regression models were used to evaluate the association of imaging metrics with known clinical diagnostic correlates such as clinical comorbidities and cardiovascular risk via Reynold’s risk score, after controlling for age and cohort. Regression t-tests were used to determine the significance of the estimated linear coefficients. The false discovery rate (FDR) was controlled for using the Benjamnini-Hochberg (BH) procedure (Benjamini & Hochberg, 1995). Significant p-values were determined using a threshold of p ≤ 0.05 after BH correction. For all multivariate regression models, regional imaging metrics were considered as the response variables.

## 3 RESULTS

Demographic characteristics are presented in **Table 1**. Characteristics are displayed for cohorts defined by HIV-status. There was no difference in age between the groups. However, there were significantly more males than females in both study populations. The HIV+ group displayed higher WMH volumes (i.e., higher CSVD burden) than controls, consistent with higher CSVD readings as shown by the DWMH and PVWMH Fazekas scores.

**Table 1.** Demographic Table. Where appropriate, values are presented as mean (standard deviation). Categorical variables are presented as number True:False. DWMH: deep white matter hyperintensity; PVWMH: periventricular white matter hyperintensity. For WMH Fazekas scores, numbers are presented as 0:>0. W:B:O corresponds to White:Black:Other/Not Reported. Hispanic numbers are presented as Hispanic:Not Hispanic:Not Reported. Sex is presented as number Female:Male; One HIV− subject identified as transgender. Units: Age (years); CD4+ Cell Count (Count/mm^3^); Viral Load (Copies/mL); Absolute WMH Volume (cm^3^).

| Category | Variable | HIV+ | HIV– | P-value |
| --- | --- | --- | --- | --- |
| **Basic** | N | 101 | 102 | 0.944 |
|  | Age | 53.20 (10.90) | 50.33 (16.72) | 0.389 |
|  | Sex | 28:73 | 32:69 | < **0.001** |
|  | Hispanic | 12:88:1 | 7:94:1 | < **0.001** |
|  | W:B:O | 52:35:14 | 83:11:8 | < **0.001** |
| **Clinical** | Diabetes Type II | 9:92 | 0:102 | < **0.001** |
|  | Hepatitis C | 11:90 | 1:101 | < **0.001** |
|  | Hypertension | 32:69 | 23:79 | < **0.001** |
|  | Reynold Risk | 6.60 (6.38) | 5.98 (6.24) | 0.088 |
| **HIV** | CD4+ Count | 672 (318) | – | – |
|  | Viral Load | 8.75 (17.7) | – | – |
| **CSVD** | DWMH | 31:70 | 43:58 | < **0.001** |
|  | PVWMH | 63:38 | 79:22 | < **0.001** |
|  | WMH Volume | 0.075 (0.71) | 0.020 (2.34) | **0.012** |

**Figure 1** shows example maps of T1w, R2*, and QSM images, with regions of interest overlaid on the anatomical image. All images are displayed in standard MNI152-2mm space. **Figure 2** shows FLAIR, R2* and QSM images of one example subject without any signs of CSVD and one example subject with both deep and periventricular WMHs present. WMH lesions are indicated in blue overlaid onto the FLAIR images. Red arrows on the QSM and R2* images indicated areas with WMHs. Blue arrows indicate NAWM. The two subjects do not appear to show localized changes in either QSM or R2* as a result of WMH lesions.

**Figure 1.**
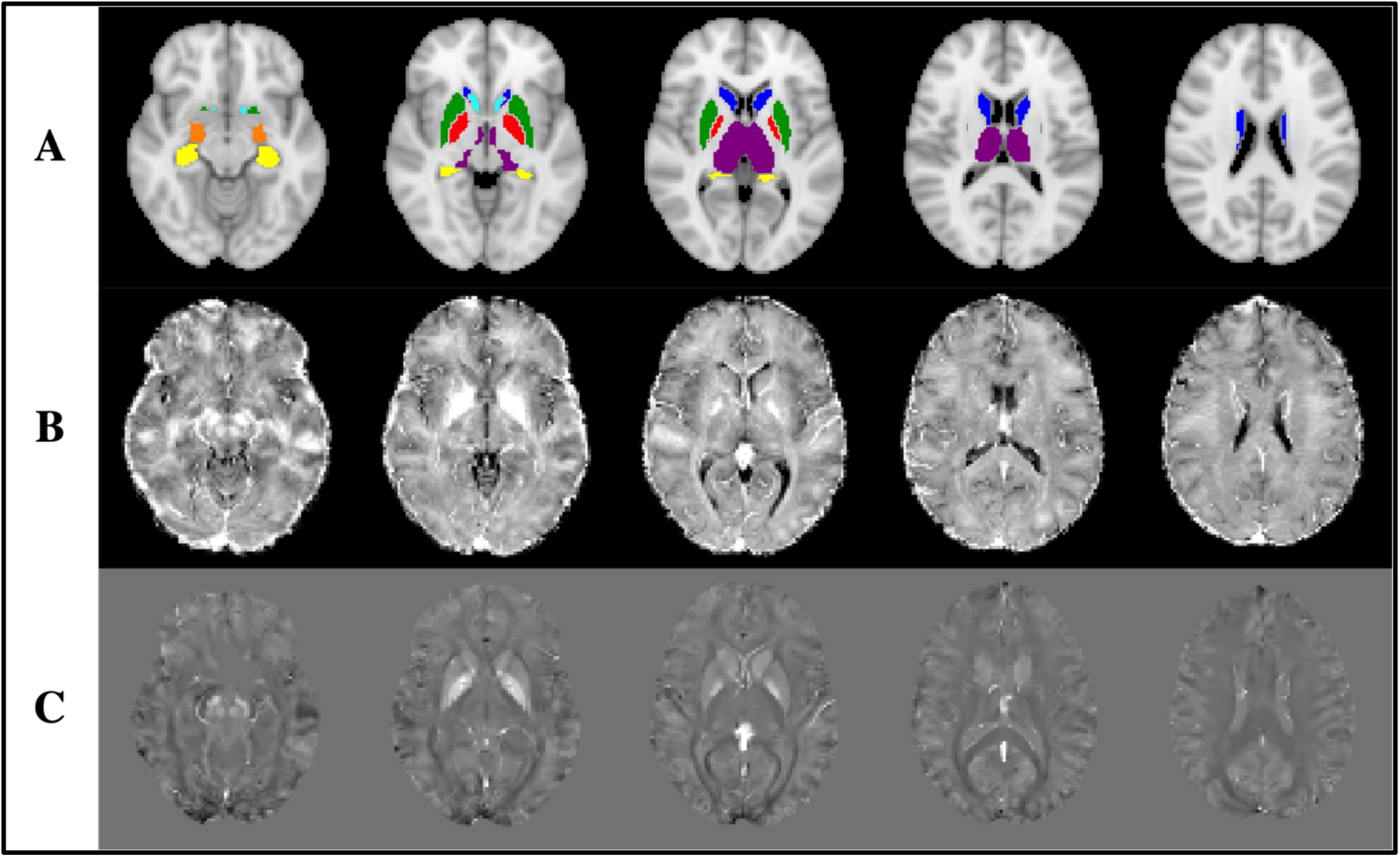
Example axial images of **A.** T1w image with subcortical regions of interests overlaid; **B.** R2* mapping; and **C.** QSM in standard MNI152-2mm space. Regions of interest include: Nuclear Accumbens (cyan), Amygdala (orange), Caudate (blue), Hippocampus (yellow), Globus Pallidus (red), Putamen (green), Thalamus (purple), Gray Matter (n.d.), White Matter (n.d.), and Whole Brain matter (n.d.). Note: n.d. – not displayed; QSM range: (−200, 200) ppb. R2* range: (0-32) s^−1^.

**Figure 2.**
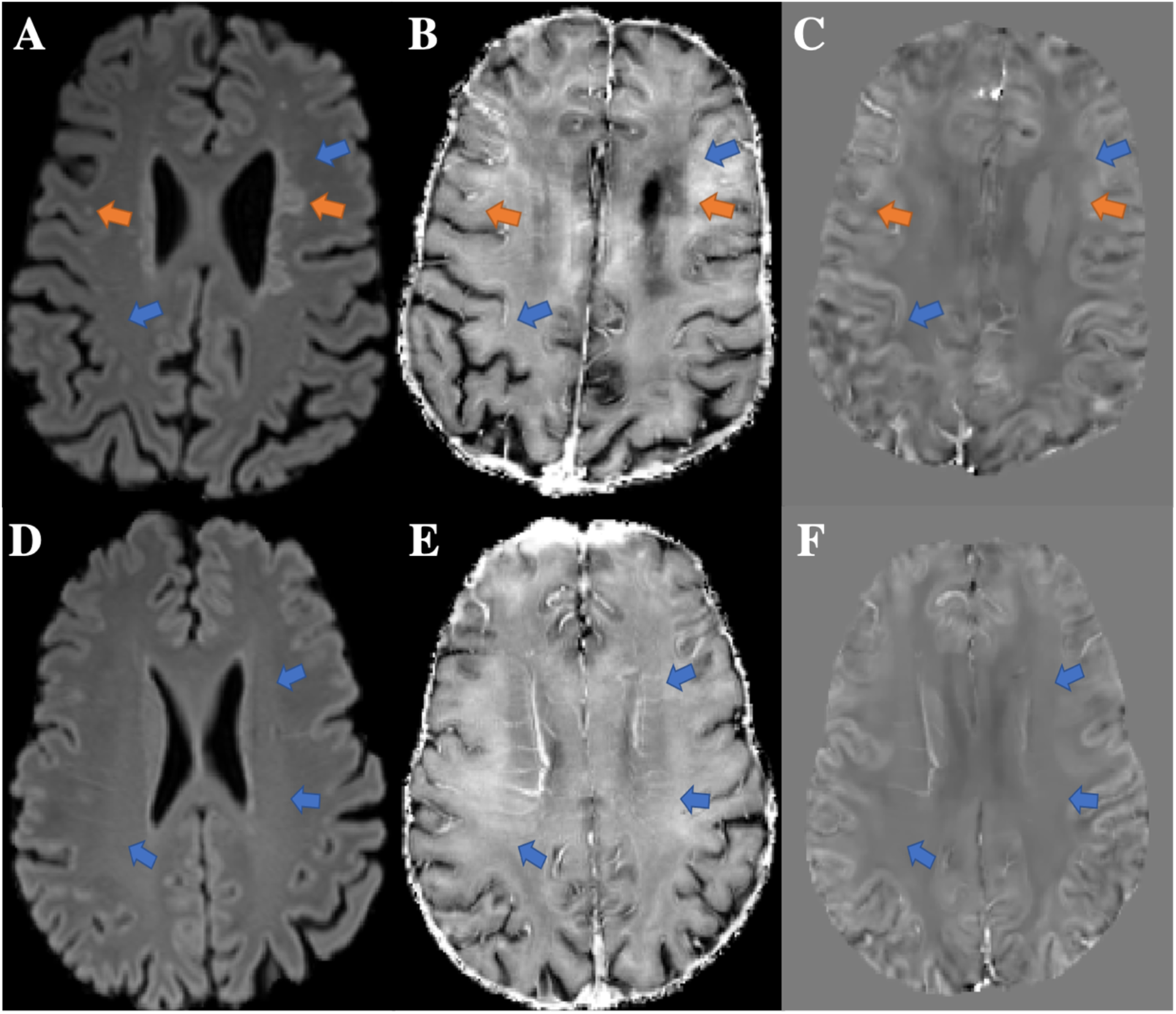
**A.** FLAIR, **B.** R2*, and **C.** QSM of one subject with white matter hyperintensities (WMHs). **D.** FLAIR, **E.** R2*, and **F.** QSM of one subject without WMHs. Orange arrows point to selected deep and periventricular WMHs. Blue arrows indicate normal appearing white matter (NAWM). Note that that R2* and QSM slices are reconstructed using a slice thickness of 2mm, which appears to show slight deviations in gyral anatomy compared to the FLAIR images. QSM range: (−200, 200) ppb. R2* range: (0-32) s^−1^.

From the HIV-status comparisons, we did not find any significant differences in QSM in any of the ROIs. QSM VBA did not show any differences between the HIV+ and HIV− cohorts. R2* VBA indicated one small potential area of significance (p < 0.05) within the posterior division of the left inferior temporal gyrus. After stratifying the HIV cohorts by CSVD status, **Table 2** shows mean QSM values and standard error for each of the four groups in each ROI. None of the ROIs showed any differences in average QSM mean values by two-way ANOVA F-tests nor post-hoc between group comparisons. While none of the between group comparisons were statistically significant, we did find at least one of two trends between group QSM values in nearly every ROI: subjects with WMHs had higher QSM than those without, independently of HIV (i.e., CSVD− < CSVD+) and subjects with HIV had lower QSM than those without, independent of CSVD (i.e., HIV+ <HIV−).

**Table 2.** Mean regional QSM values and (standard error) for each of the four cohorts {HIV− CSVD−; HIV−CSVD+; HIV+ CSVD−; and HIV+CSVD+}. No significant group differences were found using two-way ANOVA F-tests nor post-hoc group comparisons.

| ROI | HIV-CSVD- |  | HIV-CSVD+ |  | HIV+CSVD- |  | HIV+CSVD+ |  |
| --- | --- | --- | --- | --- | --- | --- | --- | --- |
| Nac | 22.18 | (2.46) | 25.83 | (1.89) | 20.73 | (2.40) | 22.84 | (1.96) |
| Amyg | -19.17 | (1.34) | -19.95 | (1.06) | -19.62 | (1.36) | -19.71 | (1.07) |
| Cau | 10.85 | (1.94) | 11.24 | (1.34) | 8.20 | (1.96) | 9.54 | (1.08) |
| Hip | -14.64 | (1.15) | -14.22 | (0.85) | -15.36 | (1.10) | -14.46 | (0.80) |
| Put | 11.33 | (2.12) | 15.83 | (1.95) | 8.90 | (2.72) | 13.25 | (1.61) |
| GPal | 50.16 | (3.49) | 62.18 | (3.42) | 53.69 | (5.57) | 61.42 | (3.01) |
| Tha | -7.02 | (0.92) | -6.48 | (0.85) | -7.49 | (1.52) | -7.05 | (0.87) |
| GM | -15.89 | (1.01) | -14.76 | (0.85) | -16.63 | (1.32) | -14.85 | (0.74) |
| WM | -17.21 | (1.05) | -16.08 | (0.89) | -18.06 | (1.42) | -16.37 | (0.76) |
| WHB | -16.38 | (1.02) | -15.23 | (0.85) | -17.13 | (1.34) | -15.40 | (0.73) |

Marginal associations between QSM and R2* and Age and HIV-specific variables are presented in the Supplementary Material. QSM moderately increased (0.15 < r < 0.39) with age in every ROI except the Cau in both cohorts. Similarly, R2* showed previously reported correlations in many subcortical ROIs (r > 0.23 in the GPal, Nac, and Put; r < −0.27 in the Cau and Tha). These results confirm that changes in tissue susceptibility distribution in the deep gray matter (DGM) are attributable to age. R2* relationships within the HIV+ cohort of HIV-specific variables showed moderate positive associations with CD4+ cell count at baseline in the GM, WM, and WHB (0.261 < r < 0.272; p = 0.022) only (**SM Table 2.1**). No other imaging metrics showed any relationships with CD4+ cell count or VL.

Marginal associations between QSM and RRS are shown in **Figure 3**. Marginal associations of QSM with age and R2* with demographic variables is displayed in the Supplementary Material. QSM increases as cardiovascular risk increases in nearly every ROI (r > 0.16) pooling all subjects. More interestingly, the within cohort correlations (HIV+ and HIV−) show that QSM is more strongly and consistently correlated with RRS in the HIV+ cohort (every ROI except the Nac), while the HIV− cohort is not significantly correlated in the Nac, Amyg, Cau, and Hip. R2* increases with RRS in the GPal (p < 0.001) and Put (p = 0.01) and decreases in the Cau (p < 0.001), pooling all subjects. These findings suggest that tissue susceptibility changes across the entire brain reflect underlying changes in iron deposition and distribution patterns due to increased cardiovascular risk.

**Figure 3.**
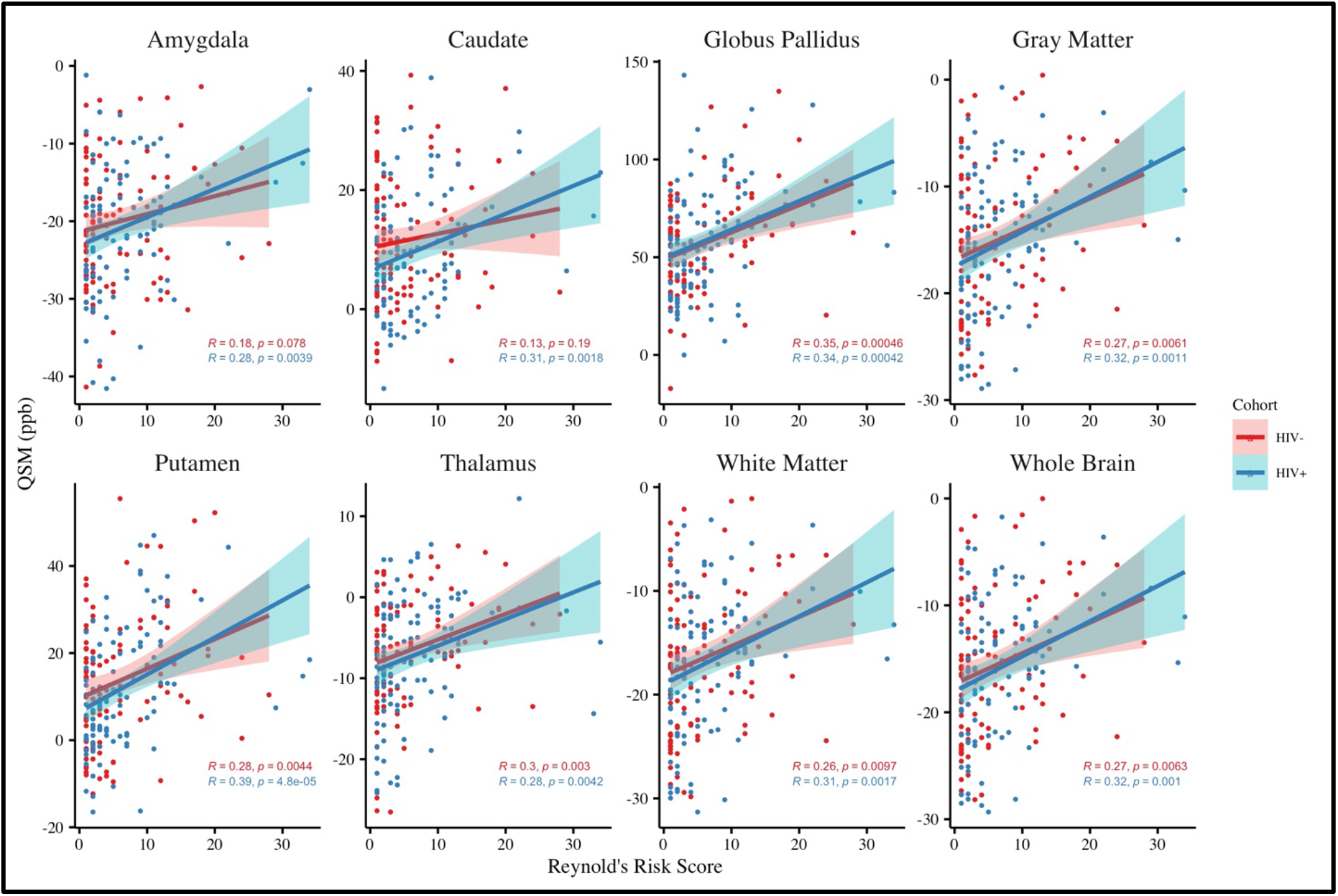
Marginal association results for regional QSM (units ppb) and Reynold’s risk score for each HIV cohort. The red and blue points represent data from the HIV− and HIV+ populations, respectively. Pearson correlation coefficients and p-values are displayed for each cohort. Regression lines are drawn with 95% confidence intervals for each cohort.

**Figure 4** shows marginal associations of demographic and regional QSM variables with absolute WMH volumes (log transformed). We found strong positive correlations between WMH volumes and cardiovascular risk (**Figure 4A**) and Age (**Figure 4B**) pooling all subjects. In the DGM, the GPal and Put showed moderate to strong associations between QSM and WMH volumes in both the HIV+ and HIV− groups, while in the Amyg and Tha, we found moderate associations between QSM and WMH volumes in the HIV+ group, only (**Figure 4C**).

**Figure 4.**
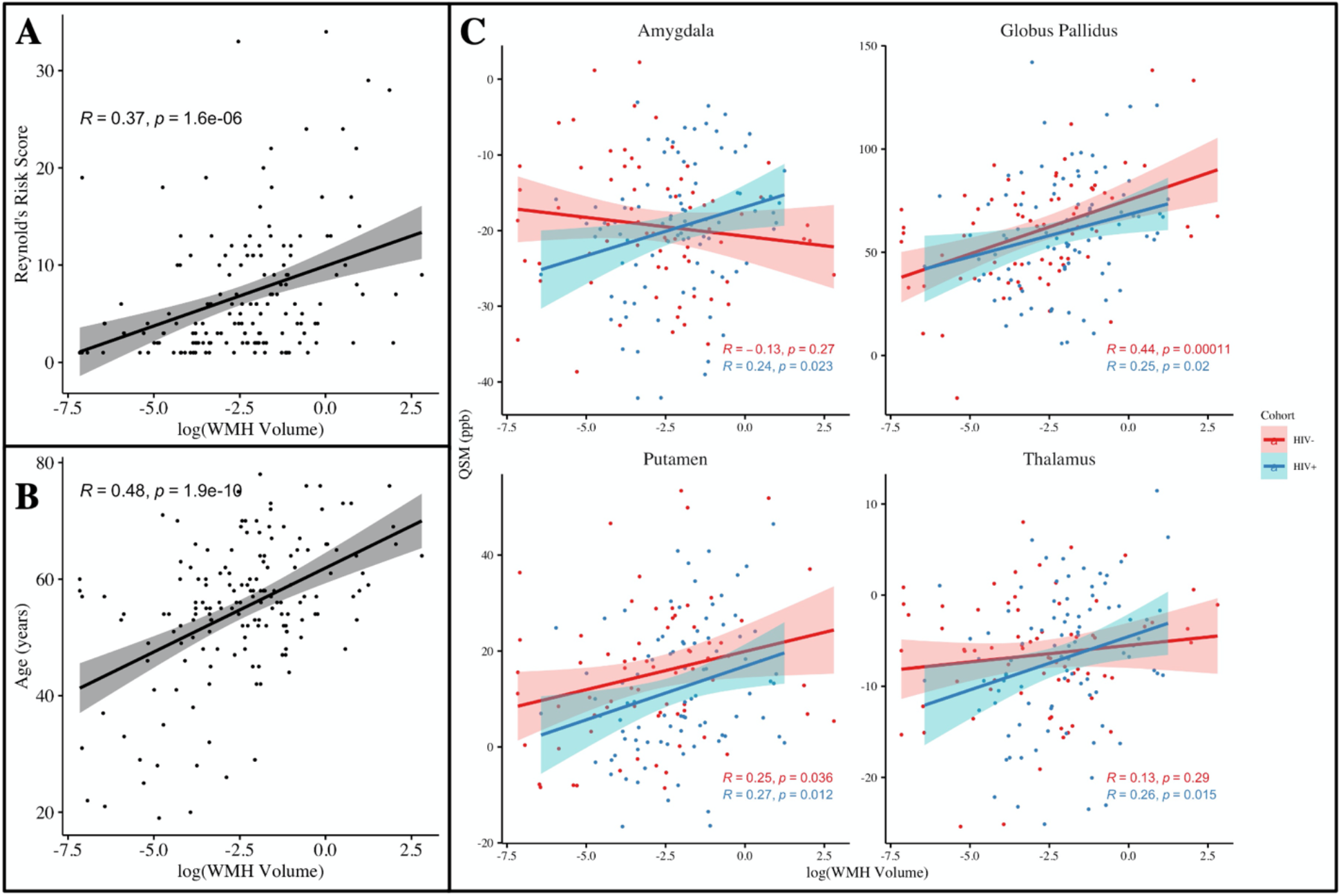
Marginal associations of **A.** Reynold’s risk score, **B.** Age, and **C.** significant regional QSM values with log transformed WMH volumes. The red and blue points represent data from the HIV− and HIV+ populations, respectively. Black points show all subjects pooled together. Regression lines with 95% confidence intervals are shown for each population. Regression lines are drawn with 95% confidence intervals for each cohort.

Associations between WMH volumes in additional regions and R2* are shown in the Supplementary Material.

Multivariate regression results are presented in **Table 3**. We found significant associations between regional QSM values and age, sex, and cardiovascular risk score. Women had higher QSM values in the Amyg and Tha and lower values in the Cau and GPal compared to men. In the Cau and GPal, after controlling for age and sex, QSM increased with cardiovascular risk score. These cardiovascular risk results confirm the relationships presented in **Figure 3**, namely that QSM increases with cardiovascular risk independent of HIV-infection. In regressions including comorbidities, we found three significant associations between imaging metrics and comorbidities before FDR correction. Specifically, QSM increases with WMH volume in the GPal (b = 2.68, p = 0.021) and R2* decreases with WMHs in the Cau (b = −0.29, p = 0.040). Finally, R2* decreases with HTN in the left and right Tha (b = −0.67, p = 0.028). None of these relationships retains significance after FDR correction.

**Table 3.** Regional QSM multivariate regression results for demographic variables: Sex, Age, Reynold’s Risk Score, and HIV-status {QSM_ROI_ ~ Age + RRS + Sex +HIV}. Regression coefficients (b – units: ppb/[variable unit]) and uncorrected p-values are displayed. Significant p-values are bolded (p < 0.05). Significant p-values after Benjamini-Hochberg correction are bolded and red (p_BH_ < 0.05).

| ROI | $b_{Age}$ | $b_{RRS}$ | $b_{Sex}$ | $b_{HIV}$ | $p_{Age}$ | $p_{RRS}$ | $p_{Sex}$ | $p_{HIV}$ |
| --- | --- | --- | --- | --- | --- | --- | --- | --- |
| Nac | 0.191 | 0.054 | 0.796 | -2.799 | 0.047 | 0.811 | 0.757 | 0.197 |
| Amyg | 0.042 | 0.226 | 3.516 | -0.047 | 0.413 | 0.061 | **0.012** | 0.688 |
| Cau | -0.102 | 0.482 | -5.794 | -1.668 | 0.108 | **0.001** | **<0.001** | 0.244 |
| Hip | 0.061 | 0.084 | 1.562 | -0.775 | 0.143 | 0.391 | 0.163 | 0.410 |
| Put | 0.268 | 0.403 | -1.942 | -2.986 | **0.001** | **0.041** | 0.389 | 0.116 |
| GPal | 0.451 | 1.035 | -10.113 | 0.311 | **0.003** | **0.004** | **0.013** | 0.927 |
| Tha | 0.069 | 0.145 | 3.318 | -0.938 | **0.101** | 0.143 | **0.004** | 0.325 |
| GM | 0.089 | 0.155 | 1.628 | -0.686 | **0.023** | 0.093 | 0.123 | 0.438 |
| WM | 0.094 | 0.145 | 1.736 | -0.876 | **0.023** | 0.132 | 0.116 | 0.344 |
| WHB | 0.088 | 0.155 | 1.679 | -0.743 | **0.025** | 0.092 | 0.112 | 0.402 |

## 4 DISCUSSION

This study evaluates and compares QSM and R2* mapping in an HIV population with increased risk for cardiovascular (CVD) and cerebral small vessel disease (CSVD) and the relationships between these MRI measures in iron rich subcortical DGM structures with cardiovascular risk scores and white matter hyperintensity (WMH) lesion volumes. The most important findings in this study were: 1) QSM and R2* did not show any significant group differences due to HIV nor CSVD as defined by a Fazekas score > 0; 2) QSM across the brain increases with cardiovascular risk independent of HIV-infection; and 3) subcortical DGM QSM increases with WMH volumes and thus CSVD burden.

Our study population shows that CSVD is more prevalent in subjects with HIV compared to controls (79% HIV + vs. 63% HIV−) in an older population as determined by higher rates and volumes of WMHs, consistent with other studies (Moulignier et al., 2018). This increased prevalence of WMHs is likely driven by chronic HIV inflammation affecting the microcirculation and vascular inflammation associated with systemic vascular risk factors (Lopez-Candales et al., 2017; Willerson & Ridker, 2004). We found significant positive correlations between the Reynold’s cardiovascular risk score and QSM in the subcortical DGM and across the entire brain in both the HIV+ and HIV− populations (**Figure 3**), which is thought to confirm that vascular risk contributes to microcirculation dysregulation resulting in increased iron deposition in the DGM. Interestingly, only the HIV+ group showed strong relationships with cardiovascular risk in the Amyg and Cau, while controls did not. In the presence of chronic HIV infection and inflammation, our results imply that a chronic inflammatory component contributes to changes in tissue susceptibility independently of the systemic vascular component.

Associations between R2* and HIV specific markers, namely positive with CD4 cell count in large ROIs (**SM Table 2.1**), may suggest that higher immune function tends to normalize R2* levels across the entire brain to that closer to controls. However, the trend of decreased 2* across the brain (**Table 2**) should be explored in future studies. For example, the exact nature of potentially decreased R2* levels may be influenced by iron dysmetabolism in mild chronic inflammation associated with HIV. Specifically, the role of microglia is altered with HIV infection and its associated immune response, which could influence tissue susceptibility throughout the brain (Cenker et al., 2017). The lack of statistical significance in individual ROIs might indicate that changes in tissue susceptibility due to immune function are not as apparent as they are when averaging over a large amount of tissue.

In other pathologies, such as multiple sclerosis (MS), neuroinflammation leads to white matter lesions that are associated with increased tissue susceptibility in and around the lesions. In the early stages of MS lesions, microglia form nodules in the NAWM without any signs of demyelination nor BBB alterations. As MS progresses, the roles of microglia change from increased activation to eventual hypocellular lesions with small amounts of microglia (Plastini et al., 2020). Throughout the MS lesion progression, QSM changes in response to the amount and roles of microglia and thus iron deposition (X. S. Zhang et al., 2019; Y. Zhang et al., 2016). The microglial response to MS lesions indicates high molecular volatility in the MS nodules. In the current study population, we found that while WMHs do not reflect tissue properties (T1w and FLAIR) of NAWM, susceptibility changes (QSM and R2*) in the WMH lesions are not different from NAWM (**Figure 2**). Since WMHs are accompanied by BBB alteration in the NAWM (Wardlaw et al., 2017), microglia are expected to be activated in the presence of CSVD. However, the lack of a change in QSM in these lesions may indicate that the role of microglia differs between MS and CSVD or that the severity of tissue injury mediated by inflammation is more severe in MS than CSVD. It should also be noted that our study population had a relatively small burden of CVSD which might have resulted in small changes in QSM.

Studies associating WMH burden and subcortical iron deposition are heterogeneous. Yan et al. (2013) demonstrates that WMHs lead to increased iron deposition via R2* in the highly vascularized subcortical DGM, while Gattringer et al. (2016) and W. Zhang et al. (2019) do not. **Table 2** supports the latter findings in that mean ROI QSM and R2* values do not statistically differ from healthy adults. However, we do see certain trends in the various groups in each region. For example, the HIV−CSVD+ mean values tend to be higher than values in the HIV− CSVD− group in every ROI except the Nac and GPal. Similarly, the HIV+CSVD+ means are higher than the HIV+CSVD− group in every ROI except the Nac and GPal. Further, **Figure 4** indicates that CSVD is more prevalent in older individuals and is associated with increased cardiovascular risk. Additionally, mean QSM in the Amyg, GPal, Put, and Tha all showed that QSM increased with WMH volume, suggesting that iron deposition in the DGM is accelerated in the presence of CSVD. The relevance of these findings requires additional investigation as they may reflect an HIV associated iron dysmetabolism mechanism. The subtle increases in QSM in the groups with WMHs and the significant regional associations with WMH volumes likely reflect changes associated with CBVD more generally.

After controlling for age, cardiovascular risk, and HIV-status, we sought to examine any effects of comorbidities associated with HIV. We found a few moderate associations between QSM/R2* and Hypertension and CSVD via the PVWMH Fazekas score and total WMH volumes. In the Cau, QSM increased with PVWMH status while R2* decreased, indicating a potential change in iron deposition and distribution in this region. Though we did not find many effects using the PVWMH Fazekas score, we did find relationships between QSM and absolute WMH volumes (**Figure 4**). Overall, QSM increased with WMH volumes independently of HIV and WMH volumes were strongly correlated with both cardiovascular risk and age. In the Tha, R2* decreased with HTN status. No other comorbidities had any significant relationships with either QSM or R2*. These results imply that comorbidities are not as influential to tissue susceptibility as age or cardiovascular risk, further confirming that QSM and R2* may only be indirectly and independently associated with HIV and CSVD.

An important implication of this work is related to dementia. Fillit et al. (2008) shows that risk factors for CVD (e.g., obesity and smoking) are associated with an increased risk of cognitive decline and dementia. QSM is known to show increased levels of brain iron deposition in dementia and related pathologies (Li et al., 2018; Moon et al., 2016). Therefore, given our relatively healthy study population, QSM may be predictive of cognitive decline and dementia as a result of CVD risk factors. Future work will include examining the relationships between QSM and cognition within this study population, since individuals with HIV are at increased risk for developing HIV-associated neurocognitive disorder (HAND) and HIV-associated dementia (HAD) (Antinori et al., 2007; Saylor et al., 2016).

Some limitations of this study include the number of males and females enrolled and an overall lack of severe CSVD burden present in both the HIV+ and HIV− groups. There is a large sex inequality in individuals with HIV; males represent a much larger proportion of newly infected individuals than females, despite both sexes having equal transmission probabilities and similar HIV risk factors, excluding lifestyle choices (Centers for Disease Control and Prevention, 2018). While our study populations do reflect similar sex demographics as normal HIV populations for both the HIV+ and HIV− groups, the female populations are underpowered compared to males. Also, in this study, subjects in both study groups show relatively small burdens of CSVD. Additionally, longitudinal changes in CSVD will be informative and will be addressed in the planned longitudinal follow-up of these subjects.

## 5 CONCLUSIONS

In this study, we investigated the relationships between the HIV and cardiovascular risk factors, and MRI-based susceptibility imaging via QSM in a cohort of HIV-infected subjects that demonstrated a higher prevalence of CSVD compared to an age-matched group of controls. Further, given the vascular and chronic inflammatory components of both HIV-infection and increased cardiovascular risk, we hypothesize that these components independently contribute to altered microcirculation, activated microglia and macrophage activity, and thus changes in iron deposition in the brain.

## Author Contributions

**Kyle Murray:** Investigation, Formal Analysis, Resources, Data Curation, Visualization, Writing- Original draft preparation. **Yi Wang, Pascal Spincemaille, Meera Singh, Md Nasir Uddin, Madalina Tivarus, Xing Qiu, Lu Wang, Henry Wang:** Data Curation, Writing-Review and Editing. **Jianhui Zhong:** Supervision, Data Curation, Writing- Review and Editing. **Sanjay Maggirwar, Giovanni Schifitto:** Conceptualization, Data Curation, Writing- Review and Editing, Funding Acquisition

## Acknowledgements

This study is funded by the National Institute of Health grant R01-AG054328. We would like to acknowledge the contributions of additional team members to the efforts of this study. Study coordinators include Jill Guary, Teresa Oh, Gillian Crysler, and Valerie Kline. Biostatistical support includes Alicia Tyrell. Schifitto imaging lab members assisting with data quality assurance and imaging support include Abrar Faiyaz, Alan Finkelstein, Arun Venkataraman, and Yuchuan Zhuang.

