## Supplementary Material for "Increased Risk for Cerebral Small Vessel Disease is Associated with Quantitative Susceptibility Mapping in HIV Infected and Uninfected Individuals"

### 1 Demographic Variable Associations

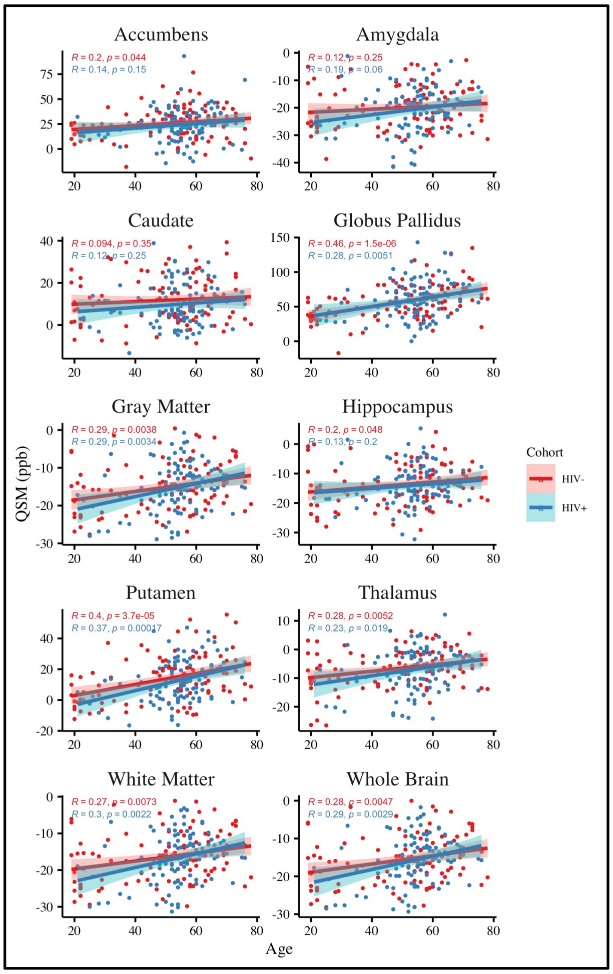

**Figure 1.1.** Marginal associations of regional mean QSM and Age for both HIV+ (blue) and HIV– (red) populations. Regression lines with 95% confidence intervals are drawn. Correlation coefficients (R) and p-values (p) are displayed for each group. QSM units: ppb. Age units: years.

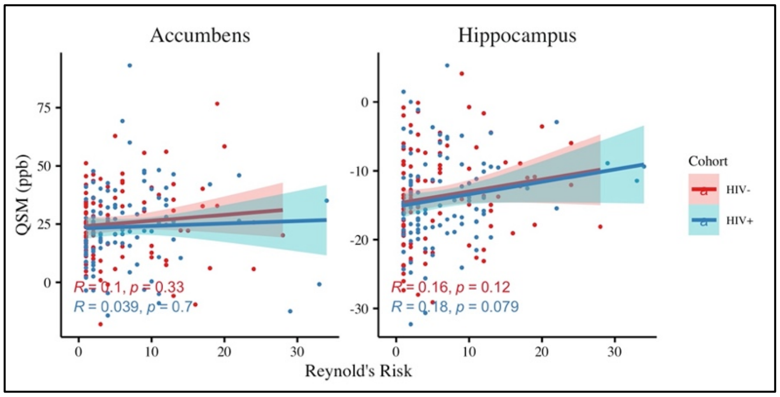

**Figure 1.2.** Marginal associations of regional mean QSM and RRS in regions where correlations are not significant. Regression lines with 95% confidence intervals are drawn.

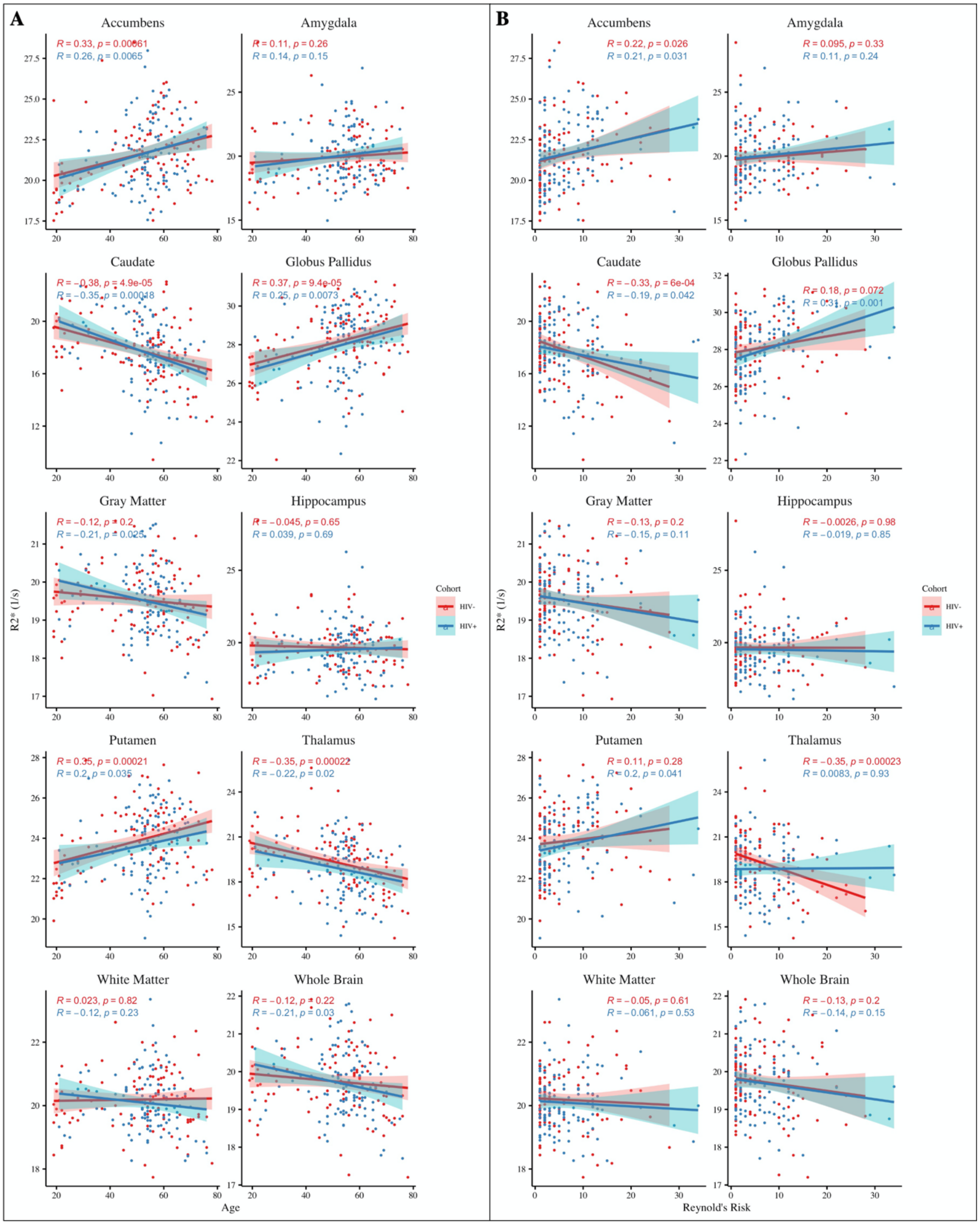

**Figure 1.3.** Marginal associations of regional mean R2* and **A.** Age and **B.** Reynold's Risk

Score for both HIV+ (blue) and HIV– (red) populations. Regression lines with 95% confidence intervals are drawn. Correlation coefficients (R) and p-values (p) are displayed for each group.

R2* units: $s^{-1}$. Age units: years.

### 2 HIV Variable Associations

| Metric | Var | ROI | R | p | p.BH | Var | ROI | R | p | p.BH |
|---|---|---|---|---|---|---|---|---|---|---|
| **QSM** | CD4 | Nac | 0.083 | 0.399 | 0.717 | VL | Nac | -0.212 | 0.029 | 0.290 |
| | | Amy | -0.070 | 0.478 | 0.717 | | Amy | -0.003 | 0.976 | 0.976 |
| | | Cau | 0.055 | 0.574 | 0.717 | | Cau | -0.016 | 0.870 | 0.976 |
| | | GPal | -0.029 | 0.768 | 0.853 | | GPal | 0.061 | 0.536 | 0.976 |
| | | Hip | -0.091 | 0.354 | 0.717 | | Hip | 0.060 | 0.544 | 0.976 |
| | | Put | -0.007 | 0.942 | 0.942 | | Put | -0.108 | 0.273 | 0.976 |
| | | Tha | -0.102 | 0.298 | 0.717 | | Tha | 0.081 | 0.410 | 0.976 |
| | | GM | -0.064 | 0.516 | 0.717 | | GM | 0.021 | 0.829 | 0.976 |
| | | WM | -0.063 | 0.523 | 0.717 | | WM | 0.010 | 0.919 | 0.976 |
| | | WHB | -0.066 | 0.499 | 0.717 | | WHB | 0.021 | 0.832 | 0.976 |
| **R2*** | CD4 | Nac | 0.166 | 0.089 | 0.165 | VL | Nac | -0.192 | 0.049 | 0.105 |
| | | Amy | 0.066 | 0.499 | 0.624 | | Amy | -0.157 | 0.107 | 0.134 |
| | | Cau | 0.166 | 0.089 | 0.165 | | Cau | -0.239 | 0.013 | 0.105 |
| | | GPal | -0.011 | 0.909 | 0.909 | | GPal | 0.006 | 0.954 | 0.954 |
| | | Hip | 0.139 | 0.157 | 0.224 | | Hip | -0.184 | 0.059 | 0.105 |
| | | Put | 0.053 | 0.587 | 0.652 | | Put | -0.175 | 0.074 | 0.105 |
| | | Tha | 0.161 | 0.099 | 0.165 | | Tha | -0.074 | 0.451 | 0.501 |
| | | GM | 0.261 | **0.007** | **0.023** | | GM | -0.217 | 0.025 | 0.105 |
| | | WM | 0.272 | **0.005** | **0.023** | | WM | -0.181 | 0.064 | 0.105 |
| | | WHB | 0.262 | **0.007** | **0.023** | | WHB | -0.205 | 0.035 | 0.105 |

**Table 2.1.** Marginal associations of regional mean imaging metrics (QSM and R2* and HIV specific variables (i.e., CD4 cell count and Viral Load (VL)) within the HIV+ cohort. Significant p-values ($p < 0.05$) after false discovery rate (FDR) correction are in bold.

### 3 R2* and WMH

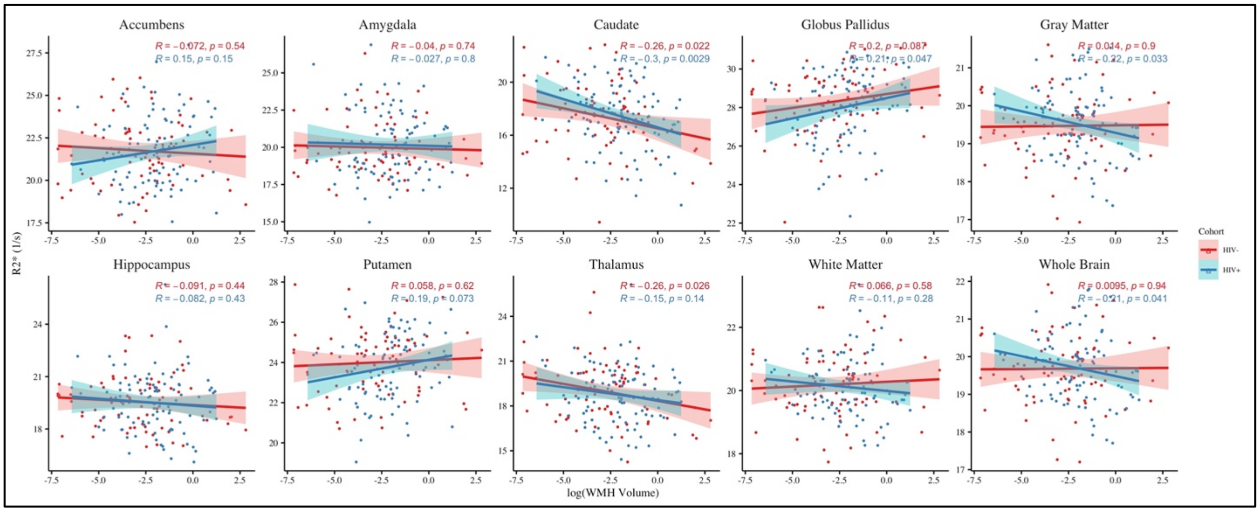

**Figure 3.1.** Marginal associations between regional mean R2* and WMH volumes (log-transformed) within the HIV+ (blue) and HIV– (red) populations. Regression lines with 95% confidence intervals are drawn. Correlation coefficients and p-values are displayed for each ROI.
